# Behavioral state regulates the dynamics of memory consolidation

**DOI:** 10.1101/2024.06.28.601231

**Authors:** N. Tatiana Silva, Maria Inês Ribeiro, Megan R. Carey

**Author notes:** **AUTHOR CONTRIBUTIONS** NTS and MRC designed the research plan. NTS and MIR performed all experiments. NTS analyzed all data and prepared figures. NTS and MRC wrote the manuscript. **DECLARATION OF INTERESTS** The authors declare no competing interests.

## Abstract

Long-term memories are consolidated over time, progressively becoming more stable and resistant to interference. Memory consolidation occurs offline and often involves transfer of memories from one brain site to another. For many motor memories, consolidation is thought to involve early learning in cerebellar cortex that is subsequently transferred to the cerebellar nuclei. Here we report that in mice, engaging in locomotor activity during training in a classical conditioning task shifts the critical time window for memory consolidation, from just after training sessions, to between trials, within sessions. This temporal shift requires natural patterns of cerebellar granule cell activity during intertrial intervals and is accompanied by earlier involvement of the downstream cerebellar nucleus. These results reveal that the critical time window for cerebellar memory consolidation can be surprisingly brief, on a timescale from seconds to minutes, and that it is dynamically regulated by behavioral state.

## INTRODUCTION

Learned associations become more stable over time through the process of memory consolidation. Consolidation has been described for a range of systems, from declarative memory to spatial memory and motor skill learning ^1–4^. Consolidation is generally thought to involve complementary cellular and systems-level mechanisms, through which specific patterns of neural activity during training, rest and sleep promote long-term plasticity and trigger a shift in memory location ^1,3,5,6^.

While often thought of as a slow process, acting across days and weeks, recent behavioral and neurophysiological studies have suggested that key aspects of memory consolidation can occur much more quickly ^7–11^. Moreover, there is evidence that consolidation may be dynamically regulated in different contexts ^12–15^.

The consolidation of motor memories is so robust that we use the term “like riding a bike” to describe things that we never forget. The cerebellum is critical for many forms of motor learning, including motor adaptation and associative learning such as classical eyeblink conditioning, in which animals learn to associate a neutral conditioned stimulus with a puff of air to the eye.Current models for cerebellar memory consolidation suggest that rapid plasticity in the cerebellar cortex leads to alterations in Purkinje cell output that then instruct further plasticity in the cerebellar nuclei ^16–19^. As a result of this process, memory storage is thought to be transferred over time, from the cerebellar cortex to the downstream nuclei ^20–23^.

Consistent with this model, pharmacologically inactivating the cerebellar cortex during a critical time window after training sessions disrupts the acquisition of cerebellum-dependent behaviors when performed early in training ^20,21,24,25^. In particular, granule cells in the cerebellar cortex have also been implicated in the consolidation process ^26^.

We recently demonstrated that cerebellum-dependent classical eyeblink conditioning is modulated by behavioral state ^27,28^. Engaging in locomotor activity enhances the acquisition and expression of conditioned responses, through mechanisms acting in the granule cell layer of the cerebellar cortex ^27^. Here we report that locomotor activity also accelerates the consolidation of eyeblink conditioning, driving a shift in the critical time window from immediately following, to within training sessions. This temporal shift is disrupted by optogenetic perturbations of granule cell activity during inter-trial intervals. It is also accompanied by earlier recruitment of the downstream cerebellar nucleus, suggesting an accelerated transfer of memory storage.

## RESULTS

We trained head-fixed mice in an eyeblink conditioning task while they were placed either on a self-paced or motorized running wheel ^27,29^. To induce eyeblink learning, a neutral conditioned stimulus (CS, here a white light LED) was paired with an unconditioned stimulus (US, here a puff of air directed at the eye) that reliably elicited an eyeblink unconditioned response (UR).

### Running makes learning insensitive to post-session inactivations of cerebellar cortex

To investigate the contribution of the cerebellar cortex (Figure 1A) in the consolidation of eyeblink conditioning in mice, we first performed reversible pharmacological inactivations of the cerebellar cortex immediately after each training session ^25,30^. We infused muscimol, a GABAa receptor agonist, into the eyelid area of the cerebellar cortex via a chronically implanted cannula (Figure 1B-D)^31–33^.

**Figure 1.**
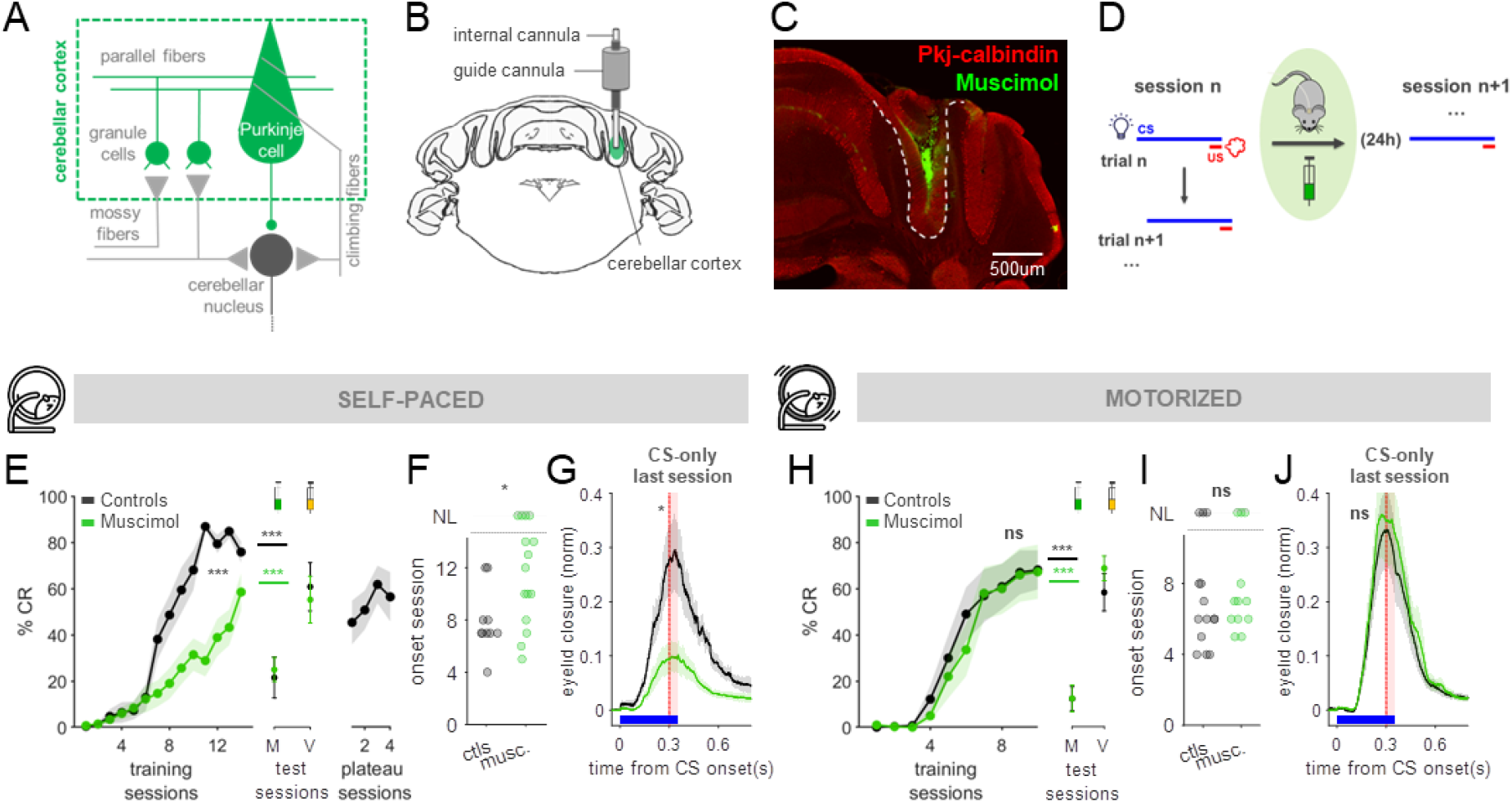
Running makes learning insensitive to post-session inactivations of cerebellar cortex. **(A)** Diagram of cerebellar cortex (green), targeted here for reversible inactivations. **(B)** Schematic of guide cannula (grey) chronically implanted at cerebellar surface and internal cannula (white) acutely introduced after each training session to deliver muscimol in the eyelid area of cerebellar cortex. **(C)** Coronal brain slice. Purkinje cells stained for calbindin (red); fluorescent muscimol (GABAa receptor agonist), green. **(D)** Protocol for post-session reversible inactivations. Muscimol or vehicle (control) was infused after each session. Learning was assessed when training resumed 24h later with the subsequent session. **(E)** Left: Averaged acquisition curves (± s.e.m.) of mice trained on a self-paced running wheel, treated with vehicle-(black, N=10) or muscimol (green, N=15) after each training session. p =0.0002***; repeated-measures-ANOVA. Center: After training completion, muscimol was infused before test sessions (M), to verify cannula placement; Controls (black): p=0.00015***; Muscimol (green): p=0.0008***; paired-t-tests, before vs. after muscimol. Performance was not impaired with vehicle infusion (V) in a subsequent session; Controls (black): p=0.5n.s; Muscimol (green): p=0.2n.s.; paired-t-tests, before vs. after vehicle. Right: Controls then received muscimol infusions after additional training sessions to test the effect of reversible inactivations after acquisition (plateau sessions, N=5). **(F)** Quantification of learning onset session (see Methods). p=0.02*; Mann-Whitney-Wilcoxon test. Non-learners (NL) did not exhibit CRs within 14 training sessions. Each dot is one mouse. (G) Averaged eyelid closures (± s.e.m.) for CS-only trials from the last training sessions. Amplitudes: p=0.046*, controls (black, N=10) vs. muscimol (green, N=15), t-test. Blue: visual CS. Red dashed line and shading: timing of the (omitted) airpuff-US. **(H)** Same as (E) but for mice trained while locomoting at a constant speed (0.1 m/s) on a motorized running wheel (Vehicle controls, black, N=14; Muscimol, green, N=12). Left: Acquisition, p=0.7n.s.; repeated-measures-ANOVA. Middle: Before vs. after Muscimol test (M). Controls: p=1.7e-6***, Muscimol, p=2.9e-6***; paired-t-tests. Right: Before vs. after Vehicle test (V). Controls, p=0.4n.s; Muscimol, p=0.1n.s., paired t-tests. **(I)** Same as (F) but for mice in (H); Controls vs. Muscimol, p=0.5n.s., Mann-Whitney-Wilcoxon test. **(J)** Same as (G) but for mice in (H); Controls vs. Muscimol, p=0.9n.s., t-test.

As previously demonstrated in rabbits ^24,25,30^, in mice walking on a self-paced treadmill, muscimol infusion in the cerebellar cortex immediately following each training session impaired acquisition of eyeblink conditioning (Figure 1E, left; compared to littermate controls infused with vehicle solution). Learning onset was delayed (Figure 1F), and the amplitudes of conditioned responses (CRs) after 14 training sessions were significantly smaller (Figure 1G). Importantly, in test sessions following training, in which muscimol or vehicle were administered before, rather than after each session, muscimol (and not vehicle) impaired CR expression equally in both training groups (Figure 1E, middle). Moreover, post-session muscimol infusions administered in control mice once learning had plateaued had no effect (Figure 1E, right), indicating that the consequences of post-session muscimol were specific to memory consolidation, and not performance or maintenance.

Intriguingly, we observed that less active mice on the self-paced wheel tended to be more susceptible than more active mice to post-session muscimol infusions (Figure S1A, open circles). To test whether locomotor activity alone could modulate the effects of post-session infusions of muscimol on memory consolidation, we externally controlled the speed at which animals walked during the training sessions ^27^. Surprisingly, we found that the same post-session infusions of muscimol in the cerebellar cortex had no effect on consolidation in mice trained on a motorized wheel rotating at a moderate fixed speed of 0.10 m/s (Figure 1H). In this case, muscimol-infused animals learned at comparable rates and reached similar CR amplitudes as vehicle controls (Figure 1H-J). This effect also appeared to be locomotor speed-dependent, as mice walking at a slower fixed speed of 0.05m/s exhibited a higher susceptibility to post-session muscimol (Figure S1B). Finally, experiments in which mice were trained on a self-paced treadmill and then transferred to the motorized wheel immediately post-session confirm that locomotor activity specifically during the training session determines susceptibility to post-session muscimol infusion (Figure S2).

These results suggest that engaging in higher levels of locomotor activity during training makes memory consolidation less susceptible to post-session inactivations of cerebellar cortex.

### Post-session optogenetic granule cell stimulation impairs memory consolidation, but only in less active states

A limitation of the pharmacological manipulations presented in Figure 1 is the difficulty of pinpointing the precise temporal window within which memory consolidation is susceptible to perturbations of cerebellar cortical activity. Although muscimol can act rapidly, its effects are likely to linger for hours ^25^. Meanwhile, cerebellar granule cells (GCs) have previously been implicated in cerebellar memory consolidation ^26^, as well as in the enhancement of eyeblink conditioning by locomotor activity ^27^. We therefore asked whether direct optogenetic perturbations of granule cell activity could reveal further insights into temporal aspects of memory consolidation and its modulation by locomotor activity.

To selectively stimulate cerebellar granule cells, we chronically implanted an optical fiber in the eyelid region of the cerebellar cortex of mice expressing ChR2 under a Gabra6 promoter ^27,29,34,35^ (Figure 2A-C). We delivered brief bursts of moderate optogenetic stimulation (5s @100Hz, every 10-15s) within a 10-minute period immediately following each training session (Figure 2D). Remarkably, in mice on a self-paced wheel, these brief bouts of post-session granule cell stimulation were sufficient to fully recapitulate the effects of cerebellar cortical muscimol infusions on memory consolidation (Figure 2E). Optogenetic stimulation during a 10-minute window after each training session delayed acquisition (Figure 2F) and reduced the amplitude of conditioned responses (Figure 2G).

**Figure 2.**
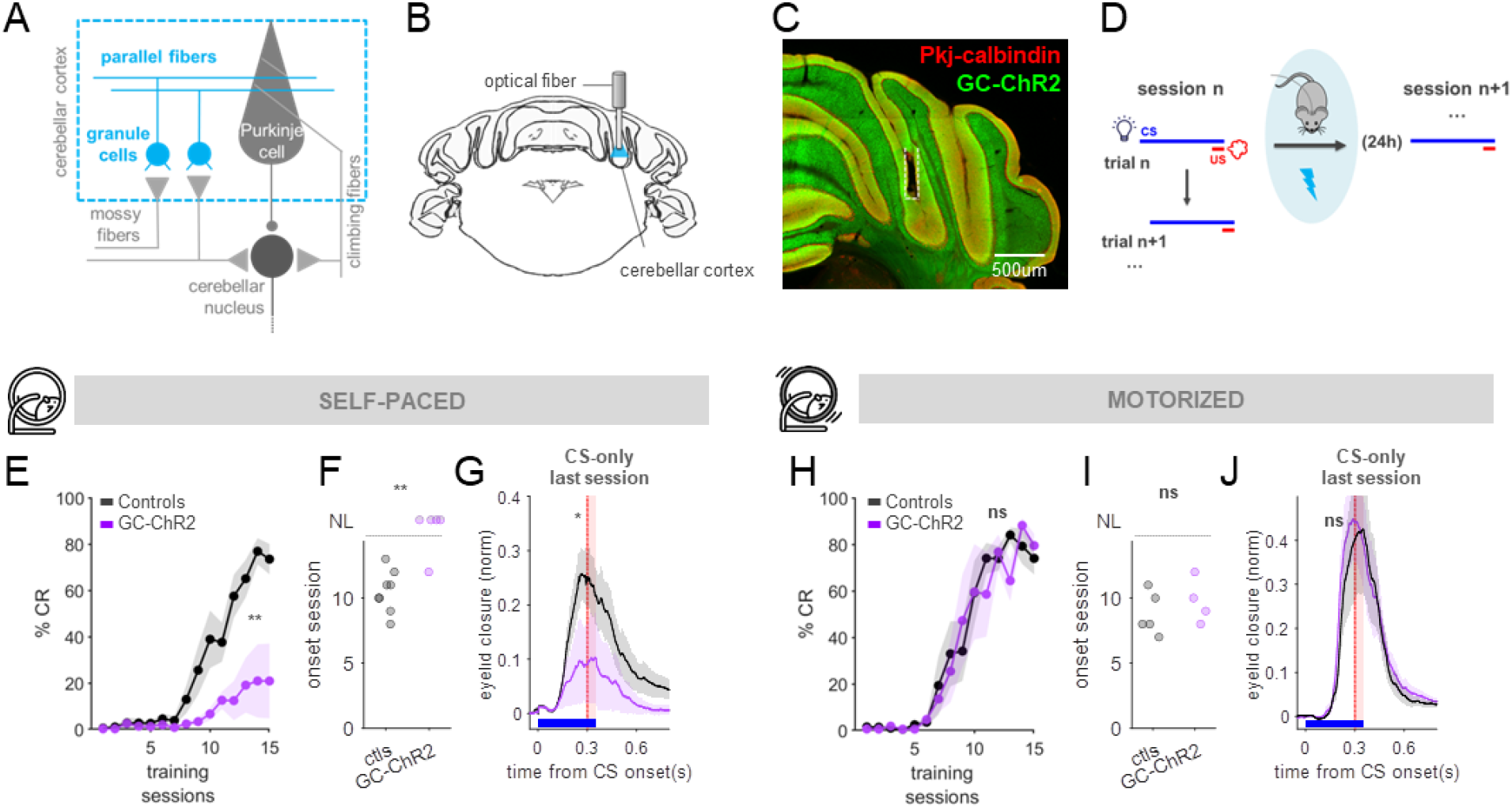
Post-session optogenetic stimulation of cerebellar granule cells impairs learning only in less active mice. **(A)** Cerebellar granule cells (blue) were targeted for optogenetic manipulation. **(B)** Schematic of optical fiber chronically implanted in the eyelid area of cerebellar cortex. **(C)** Coronal brain slice showing fiber placement in a mouse expressing ChR2 in granule cells (green). **(D)** Protocol for post-session optogenetic perturbations. Light stimulation (5s @100Hz) was presented for every 10-15s for 10-mins after each training session. Training resumed 24h later, in the following session. **(E)** Averaged learning curves of littermate controls (black, N=8) and GC-ChR2 mice (purple, N=5) trained on a self-paced running wheel. p=0.002**; Repeated-measures-ANOVA. **(F)** Quantification of learning onset session; p=0.005**, Mann-Whitney-Wilcoxon test. **(G)** Averaged eyelid closures for CS-only trials of the last training session. Amplitudes: p=0.04* t-test. **(H)** Same as (E) but for mice trained on a motorized running wheel. p=0.6n.s., Controls (black, N=5) vs. GC-ChR2 (purple, N=4); repeated-measures-ANOVA. **(I)** Same as (F) but for mice in (H); p=0.4n.s., Controls vs. GC-ChR2, Mann-Whitney-Wilcoxon test. **(J)** Same as (G) but for mice in (H); p=0.8n.s., Controls vs. GC-ChR2, t-test.

In contrast, the same post-session optogenetic stimulation of granule cells left acquisition of learning completely intact in mice that were consistently locomoting on a motorized wheel during training (Figure 2H-J). Here, as with muscimol, we also noted a similar tendency for less active animals on the self-paced wheel to be more susceptible to post-session optogenetic perturbation (Figure S1C).

Thus, engaging in higher levels of locomotor activity during training appears to protect the consolidation of eyeblink conditioning memories from susceptibility to both pharmacological and optogenetic post-session perturbations of cerebellar cortical activity.

### Engaging in locomotor activity during training accelerates the temporal window for memory consolidation

The results so far led us to hypothesize that locomotor activity may promote a shift of the critical time window for consolidation of the eyeblink learned responses, from immediately following training, to between trials, within the training session itself. To test this idea, we briefly optogenetically perturbed GCs during the intertrial intervals, within sessions (Figure 3A; Methods). Importantly, apart from being delivered within, instead of after, training sessions, the optogenetic stimulation parameters for these experiments were identical to the stimulation delivered post-session in Figure 2.

**Figure 3.**
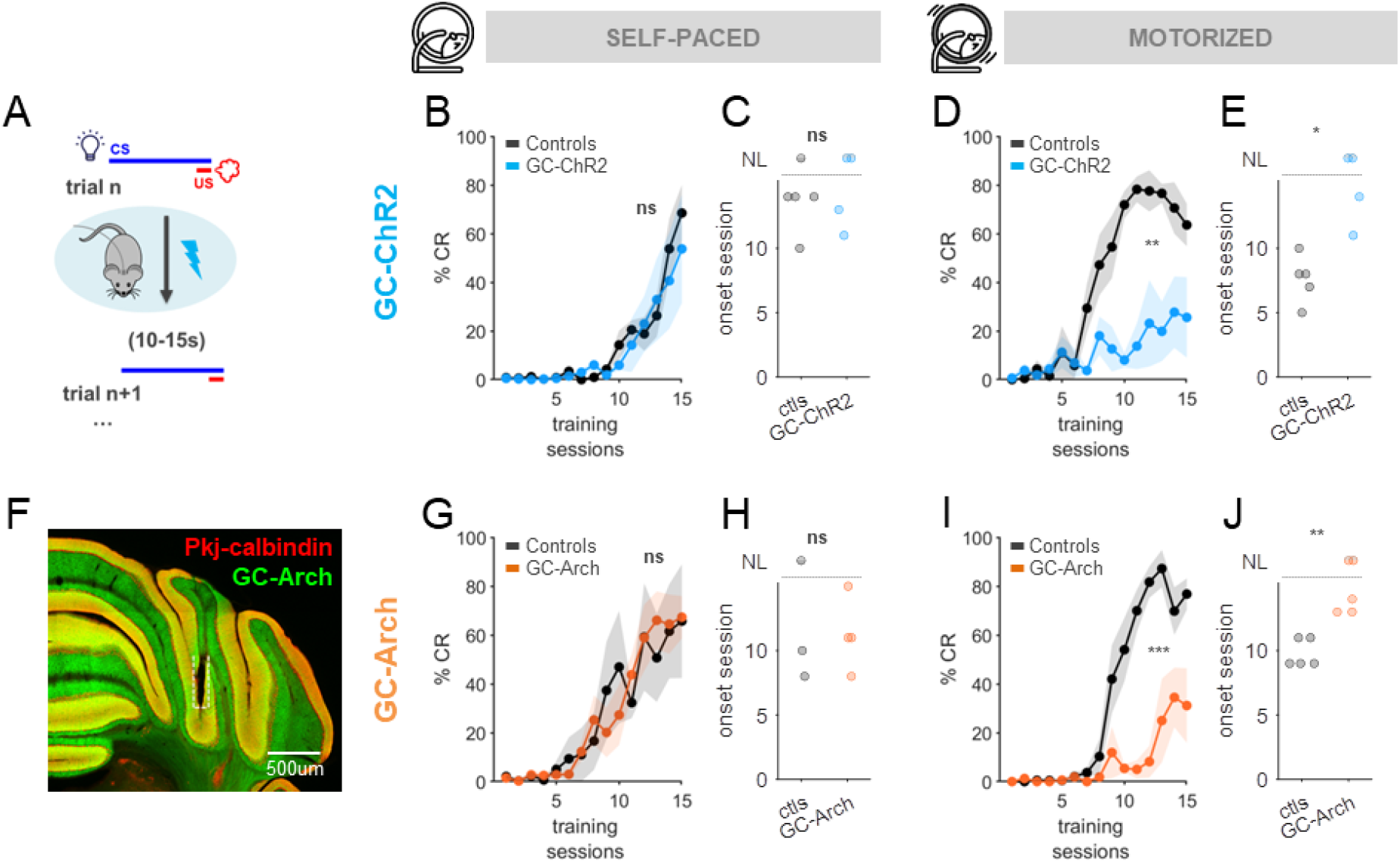
Optogenetic perturbations between trials only impair learning in mice actively engaged in locomotion. **(A)** Protocol for between-trial optogenetic perturbations of granule cells. **(B)** Averaged learning curves of littermate controls (black, N=5) and GC-ChR2 mice with ITI perturbation (blue, N=4) trained on a self-paced running wheel. Optogenetic stimulation (5s @100Hz) was delivered during intertrial intervals. p=0.7n.s., Repeated-measures-ANOVA. **(C)** Quantification of learning onset session. p=0.9n.s. Mann-Whitney-Wilcoxon test. **(D)** Same as (B) but for mice trained on a motorized wheel. p=0.002** (Controls, black, N=5 vs. GC-ChR2 with ITI perturbation, blue, N=4); repeated-measures-ANOVA. **(E)** Same as (C) but for mice in (D). p=0.02*, Mann-Whitney-Wilcoxon test. **(F)** Coronal brain slice showing fiber placement in a mouse expressing Arch in granule cells (GC-Arch). **(G-J)** Same as (B-E) but for mice expressing inhibitory Arch in granule cells. **(G)** Averaged learning curves on a self-paced treadmill. p=0.6n.s. (Littermate controls, black, N=3 vs. GC-Arch with ITI perturbation, orange, N=4); repeated-measures-ANOVA. **(H)** Same as (C) but for mice in (G); controls vs. GC-Arch, p=0.9n.s., Mann-Whitney-Wilcoxon test. **(I)** Averaged learning curves on a motorized treadmill. p=0.0003*** (Littermate controls, black, N=5 vs. GC-Arch with ITI perturbation, orange, N=5); repeated-measures-ANOVA, **(J)** Same as (C) but for mice in (I); p=0.008**, Controls vs. GC-Arch, Mann-Whitney-Wilcoxon test.

First, we optogenetically stimulated GCs between trials in mice that were trained while walking on the self-paced treadmill (Figure 3B,C). As expected, this manipulation did not affect either acquisition or expression of conditioned responses, indicating that GC stimulation itself did not substantially disrupt cerebellar encoding of the CS or otherwise directly interfere with learning. In contrast, however, identical optogenetic GC stimulation in GC-ChR2 mice running on a motorized treadmill delayed learning (Figure 3D,E). In other words, consistent with our hypothesis, granule cell stimulation between trials only impacted learning in mice that were engaged in high levels of locomotor activity – the exact opposite pattern of the results we obtained with identical optogenetic GC stimulation when it was delivered post-session (Figure 2).

To test whether the effects of granule cell perturbations within the training sessions were specific to stimulation, or whether inhibiting endogenous granule cell activity might also interfere with consolidation, we performed a similar set of experiments in mice expressing the inhibitory opsin Arch in granule cells ^36^ (Figure 3F-J). Here, too, the acquisition of learning was delayed only in the motorized treadmill condition (Figure 3I,J). Intriguingly, in both these stimulation and inhibition experiments, we also observed that animals that locomote more on the self-paced wheel tended to be *more* sensitive to optogenetic perturbations between the trials of learning (although not statistically significant; Figure S1D,E) – again, the exact opposite pattern from that observed with post-session perturbations (Figures 1 and 2).

Taken together, these results suggest that engaging in locomotor activity shifts the critical temporal window for memory consolidation from after, to during training sessions, and that this rapid consolidation requires natural patterns of cerebellar granule cell activity between trials.

### Locomotor activity accelerates the transfer of memories to the cerebellar nuclei

Cerebellar memory consolidation is thought to involve the transfer of memories from cerebellar cortex to the downstream cerebellar nuclei ^1,16,21,37^. We therefore wondered whether the temporal shift in memory consolidation we observed in mice engaged in high levels of locomotor activity might also be associated with an earlier spatial shift in memory storage.

To investigate the possibility of an accelerated shift in memory storage, we performed post-session reversible inactivation of the anterior interposed nucleus (AIP, the cerebellar nucleus for eyeblink conditioning ^17,32,38–40^), in mice on both self-paced and motorized treadmills (Figure 4A-D).

**Figure 4.**
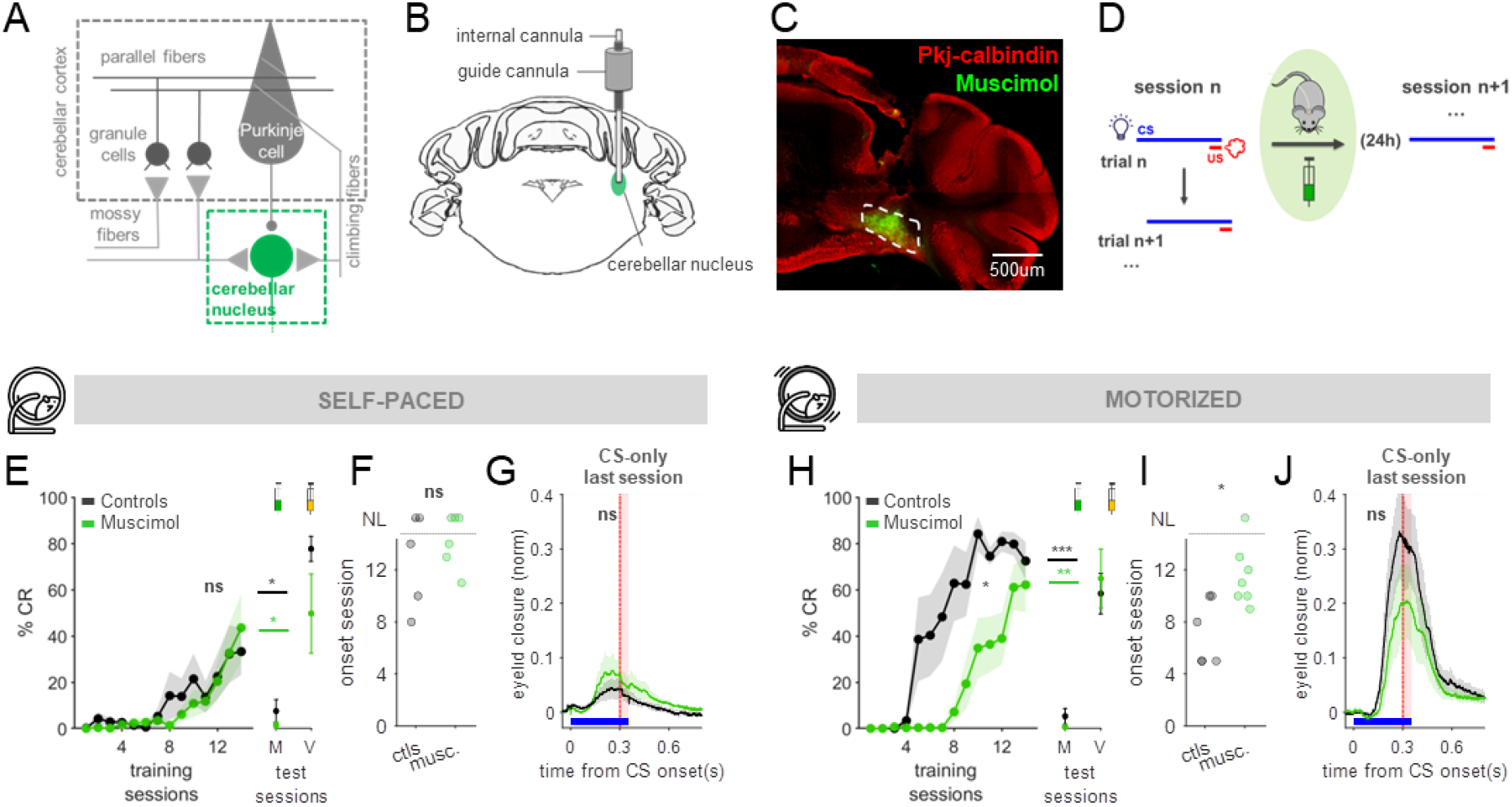
Post-session reversible inactivation of the cerebellar nucleus delays learning in mice engaged in locomotion. **(A)** The cerebellar nucleus (green square) was targeted for reversible inactivations. **(B)** Schematic of cannula implantation to infuse muscimol in the anterior interposed nucleus (AIP) cerebellar nucleus. **(C)** Coronal brain slice showing fluorescent muscimol (GABAa receptor agonist) in green. **(D)** Protocol for post-session reversible AIP inactivations. Muscimol was infused after each session and training resumed 24h later. **(E)** Left: Averaged learning curves of vehicle- (black, N=5) and AIP-muscimol-treated mice (green, N=6) trained on a self-paced wheel. p=0.6n.s., Repeated-measures-ANOVA. Right: After training, muscimol was infused before Test sessions (M), on animals in both conditions, to verify cannula placement. Before vs. after muscimol: Controls, p=0.03*; Muscimol, p=0.02*; paired-t-tests. in a following session, in a Vehicle Test session (V), performance was not impaired. Before vs. after vehicle: Controls, p=0.06n.s; Muscimol, p=0.9n.s.; paired-t-test, **(F)** Quantification of learning onset session. p=0.5n.s., Mann-Whitney-Wilcoxon test. **(G)** Averaged eyelid closure for CS-only trials from the last training session. Amplitudes: p=0.4n.s., t-test., **(H)** Same as (E) but for mice trained on a motorized wheel. Left: Acquisition, p=0.03*; vehicle Controls, black, N=6 vs. Muscimol, green, N=7; repeated-measures-ANOVA. Right: Test sessions. Before vs. after Muscimol-test (M): Controls (black), p= 0.0002***. Muscimol (green), p=0.004**. Before vs. after Vehicle-test (V): Controls, p=0.2n.s., Muscimol, p=0.8n.s. All paired t-tests. **(I)** Same as (F) but for mice in (H); p=0.01*, Mann-Whitney-Wilcoxon test. **(J)** Same as (G) but for mice in (H); p= 0.3n.s., t-test.

As expected based on previous work in rabbits ^24,30^, in mice walking on a self-paced treadmill, post-session infusions of muscimol in the AIP early in training did not affect the acquisition of learning (Figure 4E-G). In contrast, the same post-session muscimol infusions in the AIP did delay learning for mice that were actively locomoting on the motorized treadmill (Figure 4H-J; speed-modulation in Figure S1F). These results indicate that only animals engaged in higher levels of locomotor activity are susceptible to post-session inhibition of cerebellar nucleus activity early in training. This is exactly the opposite pattern from that observed with post-session perturbations of cerebellar cortex (in Figures 1 and 2), and is consistent with the hypothesis of earlier transfer of memory storage to the cerebellar nuclei in mice engaged in high levels of locomotor activity.

## DISCUSSION

We observed a double dissociation of memory susceptibility to manipulations of cerebellar activity after, versus within, training sessions, which depended on behavioral state. In relatively quiescent mice, consolidation of cerebellar memories required intact cerebellar cortical, and specifically granule cell, activity during a brief time period post-training. Engaging in higher levels of locomotor activity shifted this temporal susceptibility window to within the training sessions themselves, while also conferring memory stability against post-session perturbations of cerebellar cortex. This acceleration of memory consolidation was also associated with earlier involvement of the cerebellar nucleus. Taken together, our results reveal that cerebellar memory consolidation requires natural patterns of offline activity in the cerebellar cortex on a surprisingly rapid time scale of seconds to minutes. Further, it can be dynamically regulated by behavioral state.

### Rapid memory consolidation

Remarkably similar mechanisms have been proposed for memory consolidation across diverse memory systems ^3,4,6,15^, and a number of recent studies have suggested that key aspects of memory consolidation may take place more rapidly than previously thought ^7–11^. In particular, for human motor sequence learning tasks, behavioral and magnetic encephalographic evidence suggests that substantial memory consolidation could happen during intertrial intervals, within training sessions themselves ^7,8^.

Here, we took advantage of the temporal specificity of optogenetic perturbations ^41^ to probe the contributions of offline cerebellar cortical activity to memory consolidation at different time points. Our results demonstrate a requirement for intact patterns of cerebellar granule cell activity immediately following, or even during, training sessions, depending on task conditions. These findings suggest a possible cerebellar mechanism for rapid motor memory consolidation, operating on a time scale of seconds to minutes.

Our findings also echo classical work that has emphasized the importance of relatively brief break intervals for learning. Spacing out training periods (compared with temporally condensed, ‘massed’ training) improves memory formation across species and tasks ^42–44^. For classical conditioning like that tested here, it has been argued that intertrial interval length is a key factor for effective learning ^45^. Our results suggest a possible mechanism for this dependence: natural patterns of activity between trials during training may facilitate learning by triggering consolidation processes that have typically been thought to occur on slower time scales.

Importantly, our findings do not preclude an additional role for slower processes, such as sleep, in the consolidation process. Delay eyeblink conditioning, like many learned behaviors, emerges steadily over multiple days, and has been shown to be influenced by sleep ^15,46,47^. Instead, our results suggest that natural patterns of activity during surprisingly brief time windows during and/or immediately following training are critical for triggering the consolidation process. The full process likely still requires subsequent, slower, cellular events such as protein synthesis or structural changes, which may be sensitive to sleep.

The temporal shift in the susceptibility window for memory consolidation that we observed under more active conditions was also associated with an earlier involvement of the cerebellar nucleus in the learning process (Figure 4). Typically, it is thought that critical consolidation events must occur first in the cerebellar cortex to enable a shift in memory storage to the cerebellar nuclei.

Interestingly, however, plasticity mechanisms in the cerebellar nuclei can support surprisingly rapid learning – even within the very first training session – when activated directly ^17^. It is therefore possible that locomotor activity either accelerates, or somehow bypasses, the need for such cortical plasticity as a precursor to memory storage in the cerebellar nucleus.

### Mechanisms for dynamic regulation of memory consolidation by behavioral state

Our results reveal a critical role in the consolidation process for natural patterns of activity in the cerebellar cortex during surprisingly brief periods of seconds to minutes during and/or after training sessions. They further demonstrate that this critical temporal window for memory consolidation can be flexibly modified by task conditions – specifically, by behavioral state. How could this be achieved?

We previously found that engaging in higher levels of locomotor activity enhanced both acquisition and expression of conditioned eyeblink responses ^27^. Optogenetic circuit dissection localized that effect to the granule cell layer of the cerebellar cortex ^27^. We proposed a model whereby the multimodal integration of running-related signals and conditioned sensory stimuli in granule cells ^48,49^ would bring them closer to firing threshold and boost the CS representation at the granule cell population level ^27^. It is possible that such a mechanism of stronger parallel fiber drive during CS presentation could accelerate plasticity-related cellular signaling processes in the cerebellar cortex, nucleus, or both.

However, this working model only accounts for possible altered activity patterns *during* training trials. It doesn’t directly account for the critical role of *offline* neural activity that we observed, either after the training session (Figure 2) or, under more active conditions, during inter-trial intervals (Figure 3). Interestingly, we found that both stimulation and inhibition of granule cells during intertrial intervals disrupted consolidation on a motorized treadmill. Similarly, brief bursts of post-session granule cell stimulation were just as effective as muscimol inhibition at interfering with memory consolidation under less active conditions. Both of these findings support a requirement for natural patterns of offline activity in the cerebellar cortex during a flexible time window that is modulated by behavioral state. These surprising results are reminiscent of other, non-cerebellar memory systems, where neuronal replay has been implicated in memory consolidation ^50–54^. It is intriguing to consider the possibility that granule cells may exhibit offline pattern repetition of the engram formed during training trials, and that such replay may be modulated by behavioral state.

## METHODS

### Animals

All procedures were carried out in accordance with European Union Directive 86/609/EEC and approved by the Champalimaud Centre for the Unknown Ethics Committee and the Portuguese Direção Geral de Veterinária (Ref. Nos. 0421/000/000/2015 and 0421/000/000/2020). Mice were kept on a reversed 12-h light/12-h dark cycle with food and water ad libitum. All procedures were performed in male and female mice of approximately 12–14 weeks of age.

*Mouse lines*. WT C57BL/6J mice were obtained from The Jackson Laboratory (Jax #000664). For optogenetic experiments, as in previous studies ^27–29^ we crossed Gabra6-Cre mice, in which Cre recombinase expression was driven by the alpha6 subunit of the GABAA receptor (MMRRC 000196-UCD)^34^ with ChR2-EYFP-LoxP (Jax #012569)^36^ or Arch-GFP-LoxP (Jax #012735)^36^.

### Surgical procedures

For all surgeries, animals were anesthetized with isoflurane (4% induction and 0.5 – 1.5% for maintenance), placed in a stereotaxic frame (David Kopf Instruments, Tujunga, CA) and a custom-cut metal head plate was glued to the skull with dental cement (Super Bond – C&B). At the end of surgery, a non-steroidal anti-inflammatory painkiller (Carprofen) was administered. After all surgical procedures, mice were monitored and allowed ∼1-2 days of recovery.

*Reversible inactivations*. A 26-gauge, 5 mm length guide cannula (PlasticsOne) was implanted at the surface of the brain (RC -5.7, ML +1.9). In order to target drug infusion to the eyelid area of the cerebellar cortex (lobule IV/V/Simplex) ^31–33^, an internal cannula projecting 1.5 mm was inserted into the guide cannula acutely for each session. To target the infusions to the cerebellar nucleus (AIP) ^32,38–40^, the guide cannula was implanted 0.2 mm below the surface, and an internal projecting 1.8 mm was used.

*Optogenetic manipulations*. Optical fibers with 100μm core diameter, 0.22 NA (Doric lenses, Quebec, Canada) were lowered into the brain through small craniotomies performed with a dental drill, and positioned over the right cerebellar cortical eyelid region (RC -5.7, ML +1.9, DV -1.5) ^31–33^.

### Behavioral procedures

The experimental setup for eyeblink conditioning was based on previous work ^27,29^. For all behavioral experiments, mice were head-fixed and positioned on a Fast-Trac Activity Wheel (Bio-Serv). Depending on the experiment, the running wheel was either freely rotating by mouse impulse (self-paced) or externally maintained at a fixed speed (motorized; usually 0.10 m/s). To externally control the speed of the treadmill, a DC motor with an encoder (Maxon) was used. Mice were habituated to the behavioral setup for at least 4 days prior to training, until they walked normally at the target speed and displayed no visible distress.Locomotor activity was measured using an infra-red reflective sensor placed underneath the treadmill (PTRobotics). For Figure S1, the average speed (total distance / total time) across training sessions was used to quantify locomotor activity for each mouse; we previously showed this to be equivalent to total distance ^27,28^.

Eyelid movements of the right eye were recorded using a high-speed monochromatic camera (Genie HM640, Dalsa) to monitor a 172 × 160 pixel region at 900fps. The setup was placed inside a soundproof box kept in the dark and mice were monitored using a surveillance camera (PlayStation Eye). Lighting was provided by a small infrared light (Infaimon). Hardware was controlled and synchronized with LabVIEW or Bonsai, together with a NI PCIE-8235 frame grabber and a NI-DAQmx board (National Instruments) or a Pulse-Pal.

Most experiments consisted of three phases: Training, Test and Extinction. During Training, mice were presented with 90% CS-US paired trials and 10% CS-only trials in pseudo-randomized order. CS-only trials allowed for the analysis of the CR kinematics without the masking effect of the US-elicited reflex blink. Trials were separated by a randomized inter-trial interval (ITI) of 10-15s. At the start of each trial, the eye was monitored automatically to ensure that the eye was open for at least 1s before trial onset. In each trial, CS and US onsets were separated by a fixed interval (ISI) of 300ms and both stimuli co-terminated. The CS had 350ms duration and was a white light LED positioned ∼3cm directly in front of the mouse. The unconditioned stimulus (US) was an air-puff (40psi, 50ms) controlled by a Picospritzer (Parker) and delivered via a 27G needle positioned ∼0.5cm from the cornea of the right eye of the mouse. The direction of the air-puff was adjusted for each session of each mouse so that the unconditioned stimulus elicited a strong eyeblink. Mice performed one session per day. Each session had either 100 trials (all self-paced experiments and motorized experiments in Figure S1B) or, to maintain roughly comparable learning onsets across the two locomotor conditions, 40 trials (all other motorized wheel experiments). Test sessions were performed only for reversible inactivation experiments and are detailed below. Finally, animals performed 4 Extinction sessions consisting of 100% of CS-only trials.

Videos from each trial were analyzed offline with MATLAB (MathWorks) ^27^. The distance between eyelids was calculated for each frame by thresholding the grayscale image and extracting the minor axis of an ellipse that delineated the eye. Eyelid traces were normalized for each session, ranging from 0 (maximal opening of the eye throughout the session) to 1 (total collapse of the ellipse). Trials were classified as containing CRs if an eyelid closure occurred with normalized peak amplitude >0.1, >100ms after CS onset and before US onset.

*Littermate controls*. Because mouse behavior, and in particular locomotor and learning profiles, can vary substantially across litters, experiments were performed in multiple batches for each condition and littermates were always used as controls. For optogenetic experiments, in our previous work ^27,29^ and in pilot experiments here, there was no difference between control groups in which 1) non-opsin expressing littermate controls received laser stimulation through an implanted fiber, or 2) opsin-expressing littermate controls were implanted with a fiber but received no laser stimulation. Therefore these control conditions were pooled (Figures 2 and 3).

### Reversible inactivations

For reversible inactivations, we used muscimol (Sigma-Aldrich), a GABAa receptor agonist diluted to 1 mM in a brain-buffer vehicle solution (ACSF, Artificial Cerebro Spinal Fluid; RD Systems). At the end of each session (approx. 5min after the last trial), either muscimol or vehicle were delivered through an internal cannula lowered 1.5 mm or 1.8 mm down the guide cannula, to target the eyelid area of the cerebellar cortex or the AIP cerebellar nucleus, respectively. Infusions were 5 min at 100 nL/min (total 500 nL), with a 1ul-hamilton syringe and infusion pump (Harvard Apparatus). After infusion, 5min were allowed for substance diffusion and then a dummy cannula was placed (PlasticsOne) covering the guide cannula before putting the mouse back in the home cage to keep it clean and unclogged, and to protect the brain from possible infection.

Muscimol sensitivity and appropriate cannula placements in the eyelid regions of cerebellar cortex and AIP were evaluated during the Test phase that followed the completion of the initial Training phase ^24,25^. First, there was a Muscimol test (M), in which both Control and Muscimol training groups received a muscimol infusion, 5-10 min before a 30-trial Test session. To account for possible dips in performance due simply to effects of the testing procedure itself, on the following day mice from both groups were tested with Vehicle test session (V). Appropriate targeting was further confirmed histologically. Note that for the AIP inactivation experiments, cannula placement itself slowed acquisition for both control and experimental groups, particularly on the self-paced treadmill (Figure 4).

### Optogenetic perturbations

Light from 473 or 594 nm lasers (LRS-0473 or LRS-0594 DPSS, LaserGlow Technologies; for ChR2 excitation and Arch inhibition, respectively) was controlled with LabView. Laser light was delivered through an optogenetics patch cord (100μm core diameter, 0.22 NA) that connected to the optical fiber implanted in the animal’s brain (zirconia tip, sleeve connection).

Post-session optogenetic perturbations started 5min after the last training trial and lasted 10min. Laser stimulation was delivered in 5-ms pulses at 100 Hz for 5s, every 10-15s. Within-session laser perturbations (ChR2: 5-ms pulses at 100Hz; Arch: ON) similarly lasted for 5s and were delivered between training trials (with 10-15s inter-trial intervals).

In GC-ChR2 experiments, optic fiber placement was verified by the presence of a power-dependent blink ^27,29^ that was also used to set the stimulation intensity for each mouse. Laser power was lowered to just below the threshold for detectable eyelid movement (0.5-6 mW, avg of 3.1mW). Corresponding laser powers were used in littermate controls. Laser-evoked blink thresholds were checked in each session to confirm maintained efficacy of laser stimulation. Since GC-Arch mice did not blink in response to laser presentation (neither onset nor offset, likely due to the absence of post-inhibition rebound excitation with Arch ^55^), power was set to 6mW (max irradiance of 190.9mW/mm^2). Predicted irradiance levels were calculated using the online platform: https://web.stanford.edu/group/dlab/optogenetics. Laser powers are comparable to those of previous studies ^27,29,31^. In all cases, correct fiber placement in the eyelid area of the cerebellar cortex was confirmed histologically.

### Histology

All experiments included histological verification of infusions and fiber placement. After the experiments, animals were perfused transcardially with 4% paraformaldehyde and their brains removed. For reversible inactivation experiments, up to 1-hour before perfusion, fluorescent muscimol (ThermoFisher #M23400) was infused to verify location and spread. Brain sections (50um thick) were cut in a vibratome and stained for Purkinje cells (with chicken anti-calbindin primary antibody #214006 SYSY, and anti-chicken Alexa488 #703-545-155 or Alexa594 #703-545-155 secondary antibodies from Jackson Immunoresearch) and for cell viability (with DAPI marker). Brain sections were mounted on glass slides with mowiol mounting medium, and imaged with 5x, 10x or 20x objectives.

### Statistical analysis

Data are reported as mean ± s.e.m., and statistical analyses were performed using the Statistics toolbox in MATLAB. ANOVA, repeated measures ANOVA, Mann-Whitney-Wilcoxon and two-sample two-tailed or paired Student’s t-Tests were performed (specified in each case). Differences were considered significant at *P<0.05, **P<0.01, and ***P<0.001. No statistical methods were used to predetermine sample sizes; sample sizes are similar to those reported in previous publications ^27–29,31^. Data collection and analysis were not performed blind to the conditions of the experiments. Mice were randomly assigned to specific experimental groups without bias.

## ACKNOWLEDGEMENTS

We thank A. Machado and T. Pritchett for maintenance of mouse lines and E. Collins for technical assistance with some experiments. We thank H. Marques for the implementation of Bonsai acquisition software. We are grateful to the Carey lab, J. Jacobs, A. Joshi, and M. Orger for helpful discussions. We thank the Champalimaud Research Vivarium Staff, Histology, Microscopy and Hardware Platforms for technical support. This work was supported by a fellowship from the Portuguese Fundação para a Ciência e a Tecnologia (FCT) #BD/105949/2014 (to NTS), and grants #PTDC/MED_NEU/30890/2017 (to MRC) and European Research Council #866237 (to MRC). Additional support was provided by Congento LISBOA-01–0145-FEDER-022170, co-financed by FCT (Portugal) and Lisboa2020 under PORTUGAL2020 agreement.

## SUPPLEMENTAL FIGURES

**Figure S1.**
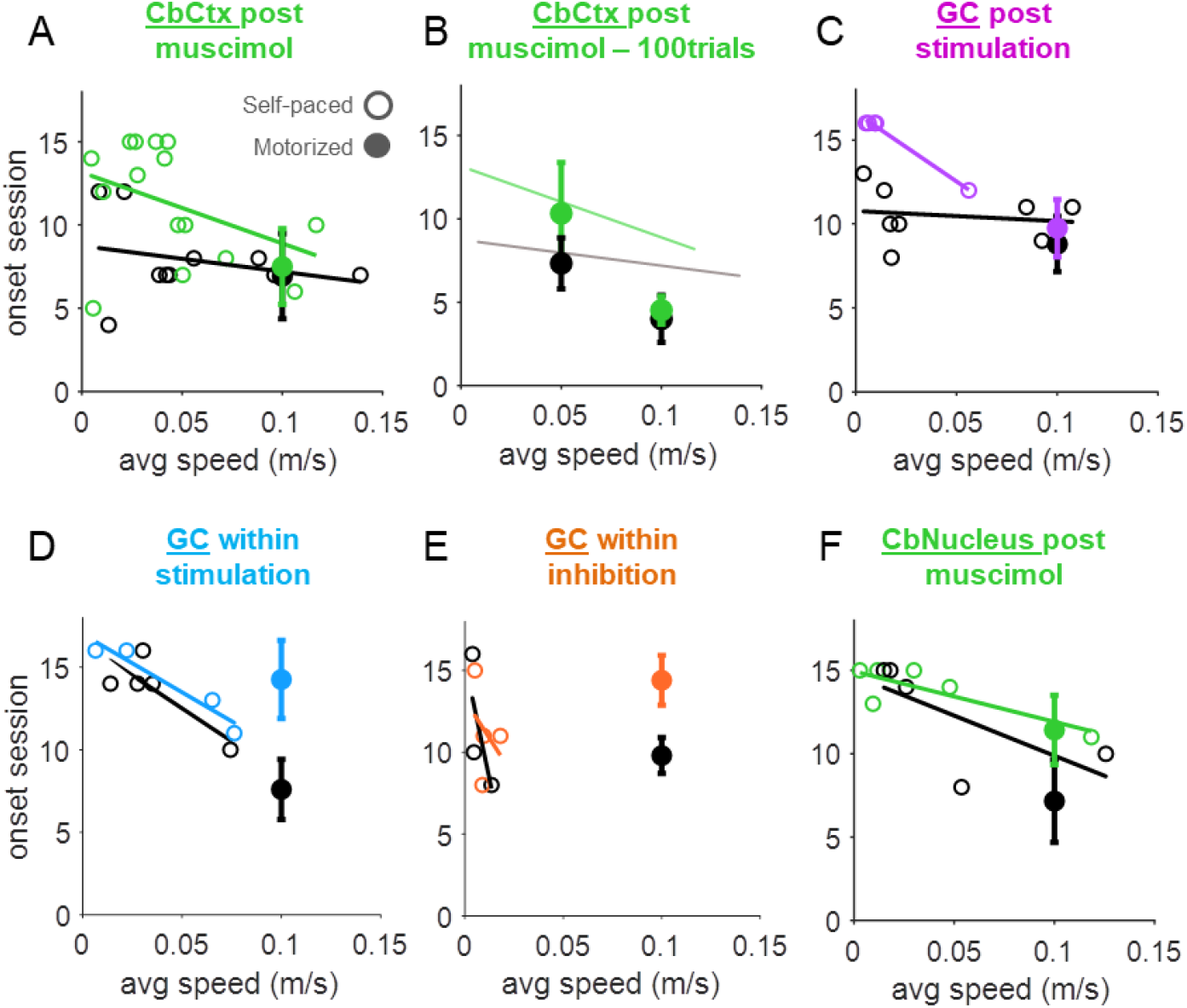
Running speed-modulation of the effects of post- and within-session perturbations on eyeblink learning. **(A-F)** Learning onset session as a function of average locomotor speed throughout training sessions, for the different groups of manipulations. Each open circle represents one mouse trained on the self-paced wheel. Closed circles represent averages (± s.e.m.) of mice trained on the motorized wheel at a fixed, constant speed. Lines represent linear regressions (y=mx+b) for the self-paced data. Controls are in black and experimental groups in color. Non-Learners were attributed an onset session value corresponding to the maximum number of training sessions for that condition +1. **(A)** Mice from the experiments in Figure 1E-G (self-paced, open circles) and H-J (motorized, closed circles). Controls (black): m=-15.5sessions/ m/s, b=8.7; Muscimol-cortex (green): m=-42.6, b=13.2. Slope: p=0.40n.s., Controls vs. Muscimol; ANOVA on linear mixed-effects model: Onsets ∼ Speeds * Conditions + (1|Mouse). **(B)** Data from mice that were trained on a motorized running wheel with longer training sessions (100 rather than 40 CS-US trials). The linear regressions from the self-paced data in (A) are re-plotted for comparison. Even on a motorized treadmill, walking at faster speeds reduces sensitivity to muscimol infusions. Onset session, 0.05m/s condition: p=0.3 n.s., Vehicle controls (black, N=3) vs. Muscimol (green, N=3); Onset session, 0.1m/s condition: p=1n.s., Vehicle controls (black, N=2) vs. Muscimol (green, N=5); Mann-Whitney-Wilcoxon tests. **(C)** Mice from the experiments in Figure 2E-G (self-paced, open circles) and H-J (motorized, closed circles). Controls (black): m=-6.03, b=10.8; GC-ChR2-post-session stimulation (purple): m=-81.9, b=16.6. Slope: p=0.03*, Controls vs. GC-ChR2, ANOVA on linear mixed-effects model. **(D)** Mice from the experiments in Figure 3B,C (self-paced, open circles) and D,E (motorized, closed circles). Controls (black): m=-79.8, b=16.5; GC-ChR2-between-trial stimulation (blue): m=-70.2, b=17. Slope: p=0.72n.s., Controls vs. GC-ChR2, ANOVA on linear mixed-effects model. **(E)** Mice from the experiments in Figure 3G,H (self-paced, open circles) and I,J (motorized, closed circles). Controls (black): m=-571.5, b=15.5; GC-Arch-between-trial stimulation (orange): m=-187.8, b=13.2. Slope: p=0.4n.s., Controls vs. GC-Arch, ANOVA on linear mixed-effects model. **(F)** Mice from the experiments in Figure 4E-G (self-paced, open circles) and H-J (motorized, closed circles). Controls (black): m=-48, b=14.7; Muscimol-AIP (green): m=-30.3, b=14.9. Slope: p=0.46n.s., Controls vs. Muscimol, ANOVA on linear mixed-effects model. Note that across conditions, mice walking at faster speeds tend to be *less* susceptible to *post*-session perturbations of cerebellar cortex (top row), but *more* susceptible to *within*-session perturbations of cerebellar cortex, or post-session perturbations of the cerebellar nucleus (bottom row).

**Figure S2.**
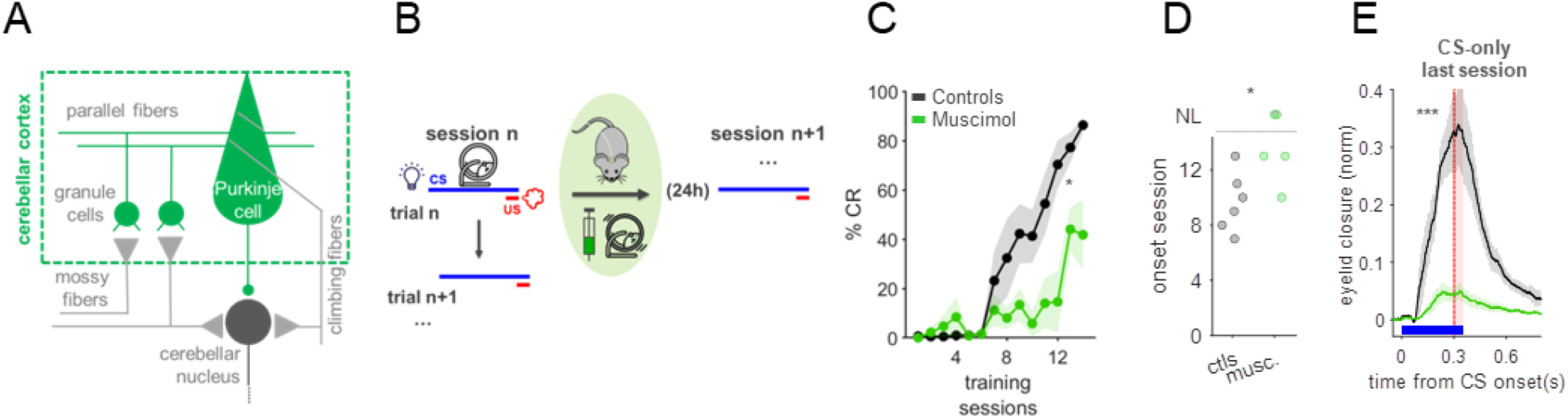
Susceptibility to post-session inactivations of cerebellar cortex is determined by locomotor activity during training. **(A)** Cerebellar cortex (green) was targeted for post-session muscimol inactivation. **(B)** Immediately following training on a self-paced wheel, mice were transferred to a motorized running wheel for muscimol infusion. **(C)** Averaged learning curves of vehicle- (black, N=6) and muscimol-treated mice (green, N=5). p=0.01*, Repeated-measures-ANOVA. **(D)** Learning onset sessions. p=0.04*, Mann-Whitney-Wilcoxon test. **(E)** Averaged eyelid closures for CS-only trials of the last training session. Amplitudes: p=0.0005***, t-test.

## REFERENCES

1. Dudai, Y. (2004). The Neurobiology of Consolidations, Or, How Stable is the Engram? Annu. Rev. Psychol. 55, 51–86. 10.1146/annurev.psych.55.090902.142050.

2. Krakauer, J.W., and Shadmehr, R. (2006). Consolidation of motor memory. Trends Neurosci. 29, 58–64. 10.1016/j.tins.2005.10.003.

3. Squire, L.R., Genzel, L., Wixted, J.T., and Morris, R.G. (2015). Memory consolidation. Cold Spring Harb. Perspect. Biol. 7, a021766.

4. Robertson, E.M., Pascual-Leone, A., and Miall, R.C. (2004). Current concepts in procedural consolidation. Nat. Rev. Neurosci. 5, 576–582. 10.1038/nrn1426.

5. Kandel, E.R., Dudai, Y., and Mayford, M.R. (2014). The Molecular and Systems Biology of Memory. Cell 157, 163–186. 10.1016/j.cell.2014.03.001.

6. Medina, J.F., Repa, J.C., Mauk, M.D., and LeDoux, J.E. (2002). Parallels between cerebellum- and amygdala-dependent conditioning. Nat. Rev. Neurosci. 3, 122–131. 10.1038/nrn728.

7. Bönstrup, M., Iturrate, I., Thompson, R., Cruciani, G., Censor, N., and Cohen, L.G. (2019). A Rapid Form of Offline Consolidation in Skill Learning. Curr. Biol. 29, 1346-1351.e4. 10.1016/j.cub.2019.02.049.

8. Bönstrup, M., Iturrate, I., Hebart, M.N., Censor, N., and Cohen, L.G. (2020). Mechanisms of offline motor learning at a microscale of seconds in large-scale crowdsourced data. npj Sci. Learn. 5, 7. 10.1038/s41539-020-0066-9.

9. Tambini, A., Ketz, N., and Davachi, L. (2010). Enhanced brain correlations during rest are related to memory for recent experiences. Neuron 65, 280–290. 10.1016/j.neuron.2010.01.001.

10. Foster, D.J., and Wilson, M.A. (2006). Reverse replay of behavioural sequences in hippocampal place cells during the awake state. Nature 440, 680–683. 10.1038/nature04587.

11. Krashes, M.J., and Waddell, S. (2008). Rapid Consolidation to a *radish* and Protein Synthesis-Dependent Long-Term Memory after Single-Session Appetitive Olfactory Conditioning in *Drosophila* J. Neurosci. 28, 3103LP–3113. 10.1523/JNEUROSCI.5333-07.2008.

12. Casagrande, M.A., Haubrich, J., Pedraza, L.K., Popik, B., Quillfeldt, J.A., and de Oliveira Alvares, L. (2018). Synaptic consolidation as a temporally variable process: Uncovering the parameters modulating its time-course. Neurobiol. Learn. Mem. 150, 42–47. 10.1016/j.nlm.2018.03.002.

13. Nguyen, N.D., Lutas, A., Amsalem, O., Fernando, J., Ahn, A.Y.-E., Hakim, R., Vergara, J., McMahon, J., Dimidschstein, J., Sabatini, B.L., et al. (2024). Cortical reactivations predict future sensory responses. Nature 625, 110–118. 10.1038/s41586-023-06810-1.

14. Tse, D., Langston, R.F., Kakeyama, M., Bethus, I., Spooner, P.A., Wood, E.R., Witter, M.P., and Morris, R.G.M. (2007). Schemas and memory consolidation. Science 316, 76–82. 10.1126/science.1135935.

15. Dudai, Y., Karni, A., and Born, J. (2015). The Consolidation and Transformation of Memory. Neuron 88, 20–32. 10.1016/j.neuron.2015.09.004.

16. Ohyama, T., and Mauk, M.D. (2001). Latent Acquisition of Timed Responses in Cerebellar Cortex. J. Neurosci. 21, 682LP–690. 10.1523/JNEUROSCI.21-02-00682.2001.

17. Broersen, R., Albergaria, C., Carulli, D., Carey, M.R., Canto, C.B., and Zeeuw, C.I. De (2023). Synaptic mechanisms for associative learning in the cerebellar nuclei. Nat. Commun., 2022.10.28.514163. 10.1101/2022.10.28.514163.

18. Bhasin, B.J., Raymond, J.L., and Goldman, M.S. (2024). Synaptic weight dynamics underlying systems consolidation of a memory. bioRxiv. 10.1101/2024.03.20.586036.

19. Boyden, E.S., Katoh, A., and Raymond, J.L. (2004). Cerebellum-dependent learning: the role of multiple plasticity mechanisms. Annu. Rev. Neurosci. 27, 581–609. 10.1146/annurev.neuro.27.070203.144238.

20. Broussard, D.M., and Kassardjian, C.D. (2004). Learning in a simple motor system. Learn. Mem. 11, 127–136.

21. Kassardjian, C.D., Tan, Y.-F., Chung, J.-Y.J., Heskin, R., Peterson, M.J., and Broussard, D.M. (2005). The Site of a Motor Memory Shifts with Consolidation. J. Neurosci. 25, 7979LP–7985. 10.1523/JNEUROSCI.2215-05.2005.

22. Shutoh, F., Ohki, M., Kitazawa, H., Itohara, S., and Nagao, S. (2006). Memory trace of motor learning shifts transsynaptically from cerebellar cortex to nuclei for consolidation. Neuroscience 139, 767–777. 10.1016/j.neuroscience.2005.12.035.

23. Kellett, D.O., Fukunaga, I., Chen-Kubota, E., Dean, P., and Yeo, C.H. (2010). Memory Consolidation in the Cerebellar Cortex. PLoS One 5, e11737.

24. Attwell, P.J.E., Cooke, S.F., and Yeo, C.H. (2002). Cerebellar Function in Consolidation of a Motor Memory. Neuron 34, 1011–1020. 10.1016/S0896-6273(02)00719-5.

25. Cooke, S.F., Attwell, P.J.E., and Yeo, C.H. (2004). Temporal Properties of Cerebellar-Dependent Memory Consolidation. J. Neurosci. 24, 2934LP–2941. 10.1523/JNEUROSCI.5505-03.2004.

26. Galliano, E., Gao, Z., Schonewille, M., Todorov, B., Simons, E., Pop, A.S., D’Angelo, E., Van Den Maagdenberg, A.M.J.M., Hoebeek, F.E., and De Zeeuw, C.I. (2013). Silencing the majority of cerebellar granule cells uncovers their essential role in motor learning and consolidation. Cell Rep. 3, 1239–1251.

27. Albergaria, C., Silva, N.T., Pritchett, D.L., and Carey, M.R. (2018). Locomotor activity modulates associative learning in mouse cerebellum. Nat. Neurosci. 21, 725–735. 10.1038/s41593-018-0129-x.

28. Albergaria, C., Silva, N.T., Darmohray, D.M., and Carey, M.R. (2020). Cannabinoids modulate associative cerebellar learning via alterations in behavioral state. Elife 9, e61821. 10.7554/eLife.61821.

29. Silva, N.T., Ramírez-Buriticá, J., Pritchett, D.L., and Carey, M.R. (2024). Climbing fibers provide essential instructive signals for associative learning. Nat. Neurosci. 10.1038/s41593-024-01594-7.

30. Attwell Rahman, S., and Yeo, C.H. (2001). Acquisition of eyeblink conditioning is critically dependent on normal function in cerebellar cortical lobule HVI. J. Neurosci. 21, 5715–5722.

31. Heiney, S.A., Kim, J., Augustine, G.J., and Medina, J.F. (2014). Precise Control of Movement Kinematics by Optogenetic Inhibition of Purkinje Cell Activity. J. Neurosci. 34, 2321LP–2330. 10.1523/JNEUROSCI.4547-13.2014.

32. Steinmetz, A.B., and Freeman, J.H. (2014). Localization of the cerebellar cortical zone mediating acquisition of eyeblink conditioning in rats. Neurobiol. Learn. Mem. 114, 148–154. 10.1016/j.nlm.2014.06.003.

33. Van Der Giessen, R.S., Koekkoek, S.K., van Dorp, S., De Gruijl, J.R., Cupido, A., Khosrovani, S., Dortland, B., Wellershaus, K., Degen, J., Deuchars, J., et al. (2008). Role of Olivary Electrical Coupling in Cerebellar Motor Learning. Neuron 58, 599–612. 10.1016/j.neuron.2008.03.016.

34. Fünfschilling, U., and Reichardt, L.F. (2002). Cre-mediated recombination in rhombic lip derivatives. genesis 33, 160–169. 10.1002/gene.10104.

35. Carey, M.R., Myoga, M.H., McDaniels, K.R., Marsicano, G., Lutz, B., Mackie, K., and Regehr, W.G. (2010). Presynaptic CB1 Receptors Regulate Synaptic Plasticity at Cerebellar Parallel Fiber Synapses. J. Neurophysiol. 105, 958–963. 10.1152/jn.00980.2010.

36. Madisen, L., Mao, T., Koch, H., Zhuo, J., Berenyi, A., Fujisawa, S., Hsu, Y.-W.A., Garcia, A.J. 3rd, Gu, X., Zanella, S., et al. (2012). A toolbox of Cre-dependent optogenetic transgenic mice for light-induced activation and silencing. Nat. Neurosci. 15, 793–802. 10.1038/nn.3078.

37. Raymond, J.L., Lisberger, S.G., and Mauk, M.D. (1996). The cerebellum: a neuronal learning machine? Science 272, 1126–1131. 10.1126/science.272.5265.1126.

38. Heiney, S.A., Wohl, M.P., Chettih, S.N., Ruffolo, L.I., and Medina, J.F. (2014). Cerebellar-Dependent Expression of Motor Learning during Eyeblink Conditioning in Head-Fixed Mice. J. Neurosci. 34, 14845LP–14853. 10.1523/JNEUROSCI.2820-14.2014.

39. Krupa, D.J., Thompson, J.K., and Thompson, R.F. (1993). Localization of a memory trace in the mammalian brain. Science (80-.). 260, 989–991.

40. ten Brinke, M.M., Heiney, S.A., Wang, X., Proietti-Onori, M., Boele, H.-J., Bakermans, J., Medina, J.F., Gao, Z., and De Zeeuw, C.I. (2017). Dynamic modulation of activity in cerebellar nuclei neurons during pavlovian eyeblink conditioning in mice. Elife 6, e28132. 10.7554/eLife.28132.

41. Goshen, I. (2014). The optogenetic revolution in memory research. Trends Neurosci. 37, 511–522. 10.1016/j.tins.2014.06.002.

42. Aziz, W., Wang, W., Kesaf, S., Mohamed, A.A., Fukazawa, Y., and Shigemoto, R. (2014). Distinct kinetics of synaptic structural plasticity, memory formation, and memory decay in massed and spaced learning. Proc. Natl. Acad. Sci. 111, E194–E202. 10.1073/pnas.1303317110.

43. Commins, S., Cunningham, L., Harvey, D., and Walsh, D. (2003). Massed but not spaced training impairs spatial memory. Behav. Brain Res. 139, 215–223. 10.1016/S0166-4328(02)00270-X.

44. Jacob, P.F., and Waddell, S. (2020). Spaced Training Forms Complementary Long-Term Memories of Opposite Valence in Drosophila. Neuron 106, 977-991.e4. 10.1016/j.neuron.2020.03.013.

45. Gallistel, C.R., and Gibbon, J. (2000). Time, rate, and conditioning. Psychol. Rev. 107, 289–344. 10.1037/0033-295x.107.2.289.

46. De Zeeuw, C.I., and Canto, C.B. (2020). Sleep deprivation directly following eyeblink-conditioning impairs memory consolidation. Neurobiol. Learn. Mem. 170, 107165. 10.1016/j.nlm.2020.107165.

47. Ramanathan, D.S., Gulati, T., and Ganguly, K. (2015). Sleep-Dependent Reactivation of Ensembles in Motor Cortex Promotes Skill Consolidation. PLOS Biol. 13, e1002263.

48. Powell, K., Mathy, A., Duguid, I., and Häusser, M. (2015). Synaptic representation of locomotion in single cerebellar granule cells. Elife 4, e07290. 10.7554/eLife.07290.

49. Ishikawa, T., Shimuta, M., and Häusser, M. (2015). Multimodal sensory integration in single cerebellar granule cells in vivo. Elife 4, e12916. 10.7554/eLife.12916.

50. Genzel, L., and Robertson, E.M. (2015). To Replay, Perchance to Consolidate. PLOS Biol. 13, e1002285.

51. Liu, X., Ramirez, S., Pang, P.T., Puryear, C.B., Govindarajan, A., Deisseroth, K., and Tonegawa, S. (2012). Optogenetic stimulation of a hippocampal engram activates fear memory recall. Nature 484, 381–385. 10.1038/nature11028.

52. Hoffman, K.L., and McNaughton, B.L. (2002). Coordinated reactivation of distributed memory traces in primate neocortex. Science 297, 2070–2073. 10.1126/science.1073538.

53. Marr, D. (1971). Simple memory: a theory for archicortex. Philos. Trans. R. Soc. London. B, Biol. Sci. 262, 23–81. 10.1098/rstb.1971.0078.

54. Diba, K., and Buzsáki, G. (2007). Forward and reverse hippocampal place-cell sequences during ripples. Nat. Neurosci. 10, 1241–1242. 10.1038/nn1961.

55. Chuong, A.S., Miri, M.L., Busskamp, V., Matthews, G.A.C., Acker, L.C., Sørensen, A.T., Young, A., Klapoetke, N.C., Henninger, M.A., Kodandaramaiah, S.B., et al. (2014). Noninvasive optical inhibition with a red-shifted microbial rhodopsin. Nat. Neurosci. 17, 1123–1129. 10.1038/nn.3752.

